# CTP synthase regulation by miR-975 controls cell proliferation and differentiation in *Drosophila melanogaster*

**DOI:** 10.1101/402024

**Authors:** Woo Wai Kan, Najat Dzaki, Ghows Azzam

## Abstract

CTP synthase (CTPsyn) is an essential metabolic enzyme. As a key regulator of the nucleotide pool, the protein has been found to be elevated in cancer models. In many organisms, CTPsyn compartmentalizes into filaments termed cytoophidia. For *D. melanogaster*, it is only its Isoform C i.e. CTPsynIsoC which forms the structure. The fruit fly’s testis is home to somatic and germline stem cells. Both micro and macro-cytoophidia are normally seen in the transit amplification regions close to its apical tip, where the stem-cell niche is located and development is at its most rapid. Here, we report that *CTPsynIsoC* overexpression causes the lengthening of cytoophidia throughout the entirety of the testicular body. A bulging apical tip is found in approximately one-third of like-genotyped males. Immunostaining shows that the cause of this tumour-like phenotype is most likely due to increased numbers of both germline cells and spermatocytes. We also report that under conditions whereby *miR-975* is overexpressed, greater incidences of the same bulged-phenotype coincides with induced upregulation of *CTPsynIsoC.* However, RT-qPCR assays reveal that either overexpression genotype provokes a differential response in expression of a number of genes concurrently associated with CTPsyn and cancer, showing that the pathways *CTPsynIsoC* affect and *miR-975* regulate may be completely independent of each other. This study presents the first instance of consequences of miRNA-asserted regulation upon *CTPsyn* in *D. melanogaster*, and further reaffirms the enzyme’s close ties to cancer and carcinogenesis.

## Introduction

One of the many hindrances to the discovery of a common cure for cancer is in the disease’s inherent complexity. Arguably every intersection of every pathway vital for fundamental cellular processes could, if disturbed, contribute to a cell’s transformation from a normal to cancerous entity (Clarke, 2017). Pathways leading to the production of nucleotides are therefore tightly regulated, as their products are not only critical for duplication and subsequent division of an emerging cancer cell (Quemeneur *et al*., 2003), but also form an integral part of energy storage (Moffatt and Ashihara, 2002) and the phospholipid-producing bio-machinery (Ostrander *et al*., 1998).

In pyrimidine synthesis, an example of an essential metabolic enzyme is CTP Synthase (CTPsyn). The enzyme catalyses the conversion of UTP to CTP via well-characterized *de novo* and salvage pathways (Kammen and Hurlbert, 1959; Lieberman, 1956). Its uniqueness lies in its tendency to form filaments called cytoophidia (Liu, 2011); such structures have been reported in all manners of organisms, ranging from the simple bacterium to advanced mammals (Carcamo *et al*., 2011; Chen *et al*., 2011; Liu, 2010; Noree *et al*., 2010). Despite their kingdom-transcending conservation, an understanding of the exact function of CTPsyn-cytoophidia as well as the factors governing their formation remains poorly established.

Certain properties of the CTPsyn-cytoophidia within *Drosophila melanogaster* has nonetheless been revealed. For example, only isoform C of its CTPsyn (CTPsynIsoC) is actually involved in the structure’s formation (Azzam and Liu, 2013). Several studies have shown a correlative relationship between cellular concentrations of the protein to the length of cytoophidia (Azzam and Liu, 2013; Noree *et al*., 2014). Under conditions whereby CTPsyn is excessively available, the structure tends to become elongated (Chen *et al*., 2011), whereas the opposite is true if its gene is knocked-out or mutated. This dynamism exhibited by the cytoophidium in response to surrounding changes means that it could, at the very least, serve as indicator for normal levels of cellular nucleotide production, especially in tissues where it could be found consistently and in relatively high numbers. The drosophilid testis is one such tissue. Both macro and micro-cytoophidia are observable in primary spermatocytes (Liu, 2010). Unfortunately, the distribution patterns of cytoophidia at points of spermatogenesis preceding and succeeding this stage are insufficiently characterized. The structure has not been reported in the germline or cyst stem cells surrounding the apical hub cells, nor the gonialblasts giving rise to spermatocytes. Whether cytoophidia is retained throughout ensuing meiotic events is yet another uncertainty.

A careful record of the trend of cytoophidia formation within testis will thus inadvertently provide information regarding CTPsyn levels at a specific point of the tissue’s development. Thorough characterization of its gene’s expression patterns is of interest as increased levels of the protein has been linked to several mammalian cancer types such as sarcoma (Weber *et al*., 1983), hepatoma (Kizaki *et al*., 1980; Williams *et al*., 1978) and leukaemia (van den Berg *et al*., 1993). Cytoophidia-forming properties of *CTPsyn* is also notably often associated with tumorigenetic genes. The popular oncogene *Myc* encourages cytoophidia formation, but requires CTPsyn to exert its cellular overgrowth effects (Aughey, Grice, and Liu, 2016). Under dysregulation, *Ack1* is linked to a poor prognosis in prostate cancer (Mahajan *et al*., 2007); CTPsyn not only colocalizes in cytoophidia with its protein, but mutants lack total CTP content, indicating that CTPsyn function is directly responsive to Ack1 (Strochlic *et al*., 2014). IMPDH, instrumental in conferring chemo-resistance to human osteosarcoma when excessively expressed, is yet another cytoophidium-forming metabolic enzyme with co-localization as well as gene-gene interaction links to CTPsyn (Fellenberg *et al*., 2010). A relationship between CTPsyn and the exacerbation of cancers is in fact observed frequently enough that it has been recognized as a feasible target in therapy. Multiple drugs have been developed with the intention of attenuating the enzyme adequately enough to combat cancer progression (Fijolek, Hofer, and Thelander, 2007; McCluskey *et al*., 2016; Verschuur *et al*., 2000).

Here we report that when the enzyme is present in excess, it contributes to an increase in the girth of the *Drosophila* testis at its apical tip. This phenotype, herewith referred to as ‘bulging’, always coincides with detection of grossly elongated cytoophidia, a hallmark of *CTPsynIsoC* overexpression in *D. melanogaster*. More detailed immunostaining reveals that bulging may be attributed to larger population sizes of germline cells and spermatocytes when compared to controls. We also show that the overexpression of a singular microRNA i.e. *miR-975* leads to similar testicular bulging phenotypes. Though it is shown that excessive growth may also be caused by increased cell numbers, and RT-qPCR data confirms the consequential upregulation of *CTPsynIsoC* in testes tissue overexpressing *miR-975*, the lengthening of cytoophidia is not observed. Our study therefore shows that whilst bulge-induction under the overexpression of *CTPsynIsoC* and *miR-975* are both related to higher levels of the CTPsynIsoC enzyme, pathways effected in either case differ greatly, inferring that CTPsynIsoC may be involved in more mechanisms of cellular growth control than previously thought.

## Results

### Testes of flies overexpressing *CTPsynIsoC* show elongated cytoophidia and significantly greater incidence of apical testis bulging (tumour-like phenotype)

We first identified the variations in phenotypes that *CTPsynIsoC* would cause in testicular tissue when it is induced to be overtly expressed under the influence of a strong, ubiquitous *Act5c* driver. All males of selective phenotype were dissected. Testes were quickly observed under the microscope. ‘Bulging’ was defined as a visibly significant increase in circumference across the apical tip when compared to the control genotype (*Act5c-GAL4 > Oregon-R*). A schematic representation of a normal testis and early spermatogenesis is represented in Figure 1A. About 34% (n=779, Figure 1B) of *Act5c-GAL4 > UAS-CTPsynIsoC* testes were overgrown (Figure 2A vs Figure 2B, Figure 2C vs Figure 2D). To show that this bulging phenotype could indeed be related to the overexpression of *CTPsynIsoC*, fixed testes were stained against CTPsyn. Cytoophidia morphology and localization patterns seen in controls (*Act5c-GAL4 > Oregon-R*) were consistent to descriptions in a previous report (Liu, 2010). Both micro and macro-cytoophidia were present in these tissues. Whereas greater concentrations of micro-cytoophidia were observed towards the end of the transit amplification (TA) region, macro-cytoophidia are often found in a more random, dispersed pattern across the same foci. No cytoophidia were detected beyond this region. Conversely, not only do bulged testes from the transgenic males (*Act5c-GAL4 > UAS-CTPsynIsoC*) show lengthened cytoophidia across the apical tips, the structure is preserved in cells of later stages of spermatogenesis as well (Figure 2E vs Figure 2F). The correlative incidence percentage between testicular bulging and cytoophidia elongation in these cases was 100%.

**Figure 1:**
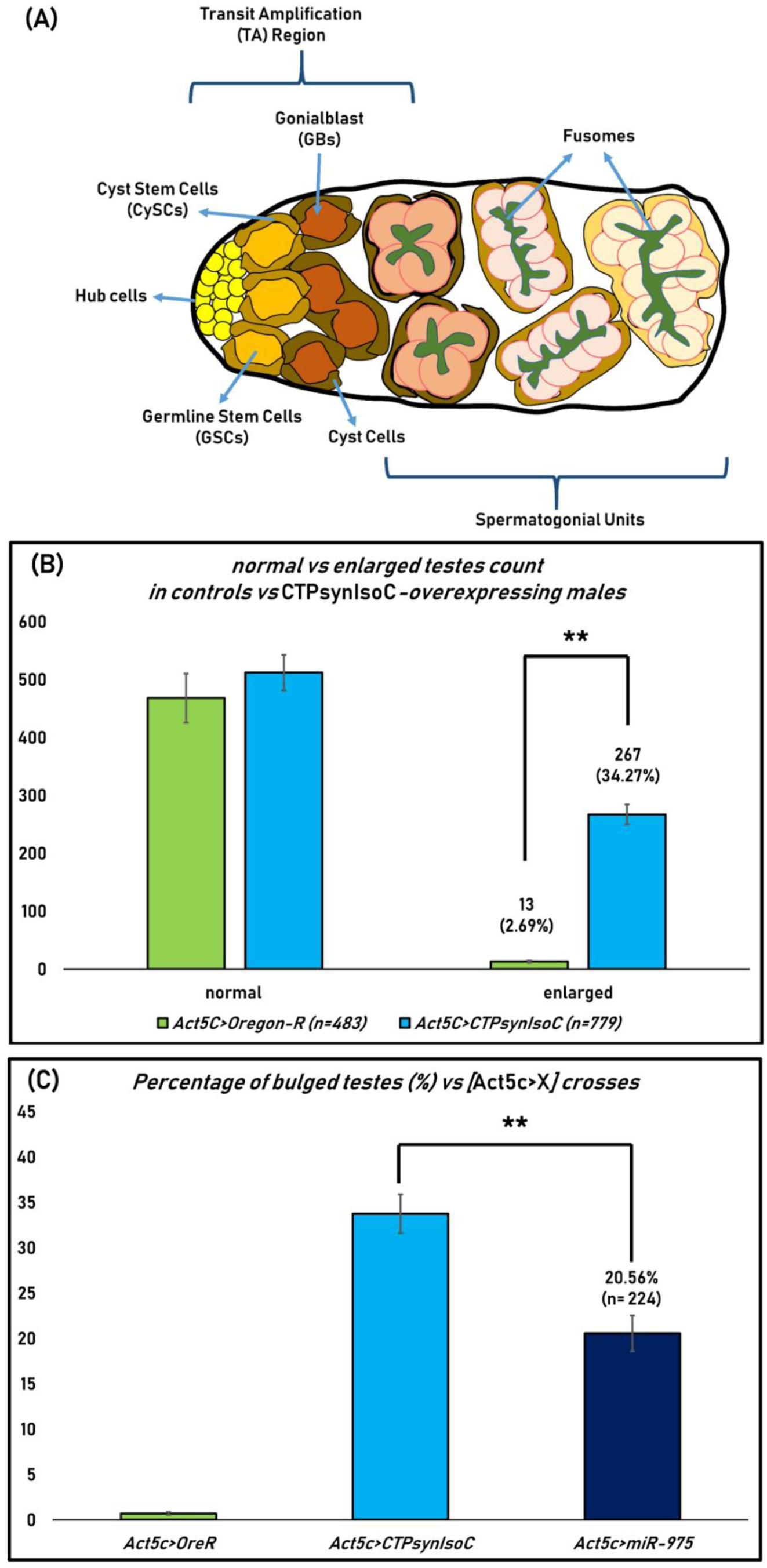
(A) Schematic diagram of spermatogenesis as it occurs from the apical tip of the testis. (B) Bar chart showing the numbers of phenotypically normal and enlarged (or bulging) testes. F1s of Oregon-R flies crossed to *w*; Act5C-GAL4 / CyO* were used as control. (C) Bar chart showing the percentage of bulged testes against *Act5C-GAL4* driver crosses. ‘**’= *p*<0.05.

**Figure 2:**
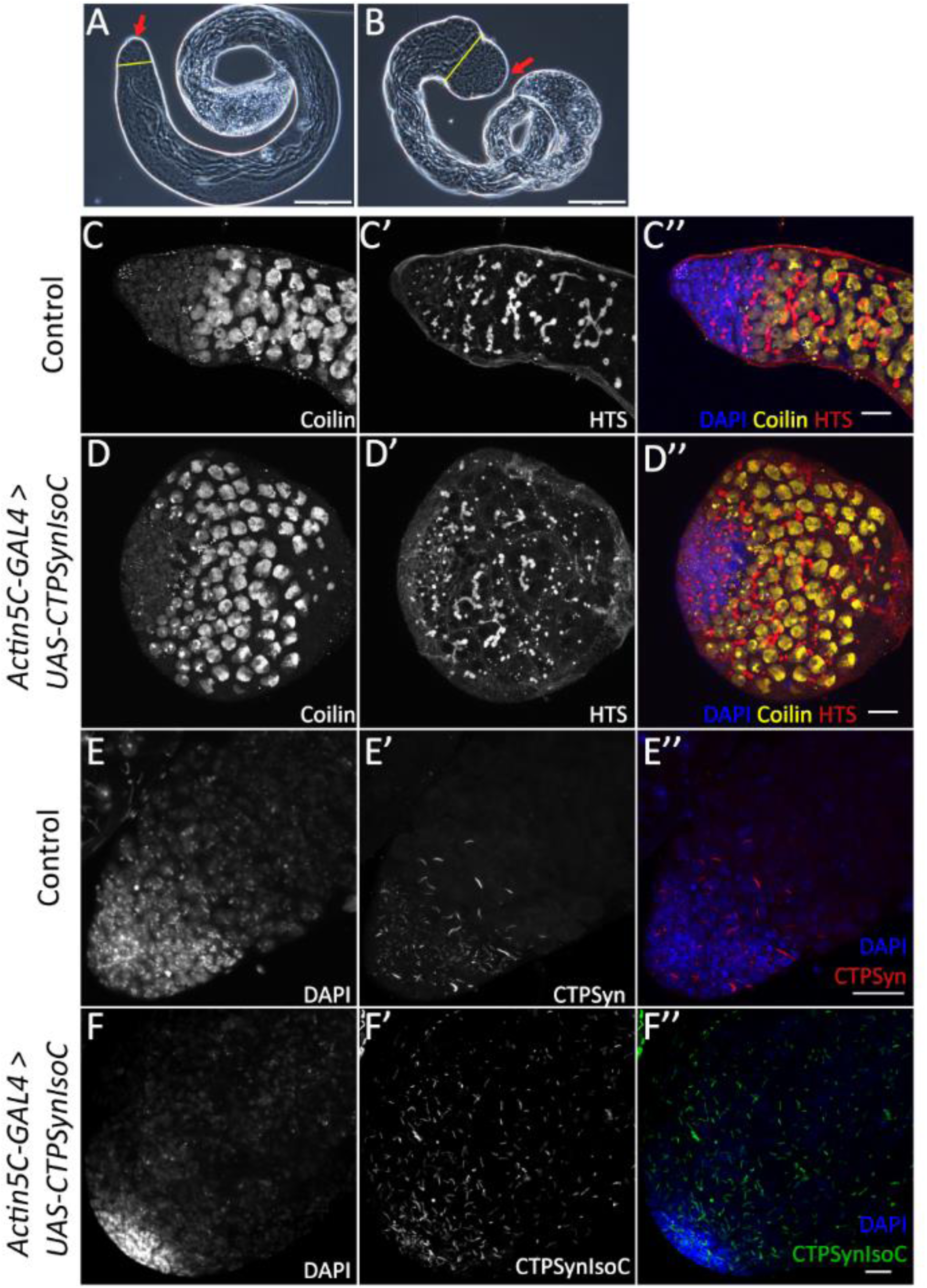
Diagrams showing control testis and transgenic testis. (A) Phase contrast image of a control testis. (B) Phase contrast image of *Act5C-GAL4 > UAS-CTPsynIsoC* testis showing the tumour-like phenotype. Red arrow indicates apical tips of the testes. Yellow line shows the relative diameter of the testis body. Scale bar: 200*μ*m. (C – C″ and E – E″) Confocal images showing the control testis. (D – D″ and F – F″) Confocal images showing CTPsynIsoC overexpression testis causing the tumour-like phenotype. Scale bar: 20*μ*m.

### Apical testes bulging arose from increased germline/spermatocyte numbers

We next attempted to identify the cause of testicular bulging through immunohistochemistry methods, refined to stain against certain markers of *Drosophila* germline stem cell (GSC) and cyst stem cell (CySC) progression. These two cell-types are bound in to the somatic hub cells (stained by Anti-FascIII) in a rosette-formation. Together, this point in the apical region of the testis is known as the stem cell niche. GSCs and CySCs divide to give rise to spermatogonia and spermatogonium-enveloping cyst cells, respectively (Figure 1A). Anti-Vasa highlights GSCs, whereas Anti-Tj (Traffic-Jam) stains both CySCs and their immediate descendants. An increased number of Vasa-positive cells was observed in *Act5c-GAL4 > UAS-CTPsynIsoC* testes compared to controls, suggesting CTPsyn-influenced growth in germline cell population (Figures 3A and 3C vs Figures 3B and 3D). GSCs themselves were however not affected by the overexpressed protein, and remained as an eight-cell rosette as is typically seen in controls (Figure 3B). An increase in Vasa-positive cells should be accompanied by an increase in Tj-positive cells, since the relationship of germline cells are interrelated to its somatic cyst cells. The opposite was instead shown to be true: in control testes, Tj-positive cells appeared to be loosely spaced, with the distance between the first and final Tj-positive cell observed extending out of the apical tip found to be longer. In *Act5c-GAL4 > UAS-CTPsynIsoC* flies, however, such cells are found closely clustered at the apical tip (Figures 4A and 4B). This suggests that the development of cyst cells is being accelerated together with the germline cells.

**Figure 3:**
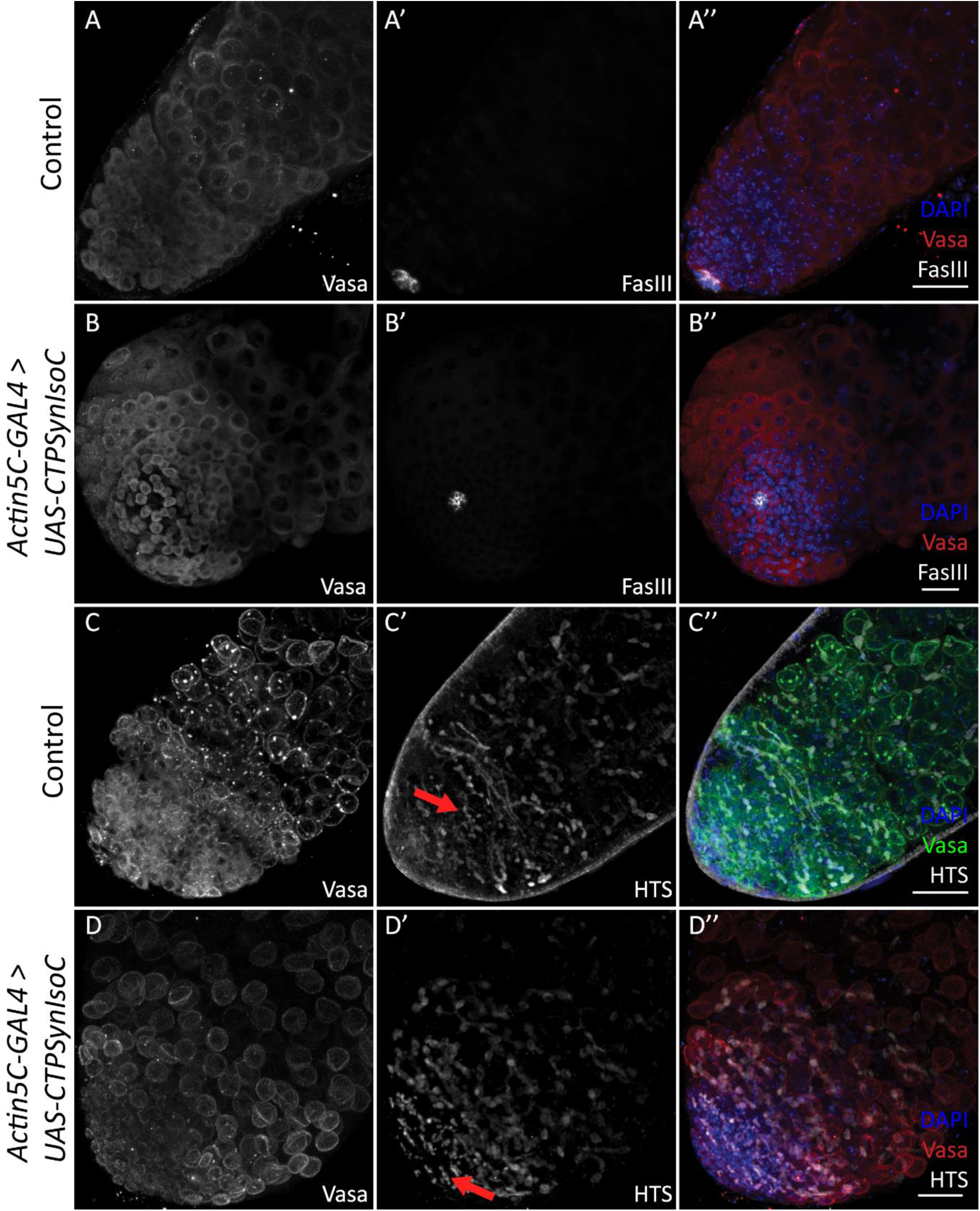
Testes stained with Anti-Vasa, Anti-FascIII, and Anti-HTS. (A-A″) Control testis stained with Anti-Vasa and Anti-FascIII. (B-B″) *Act5C-GAL4 > UAS-CTPsynIsoC* testis stained with Anti-Vasa and Anti-FascIII. Note that the numbers of GSC adjacent to the hub cells are phenotypically normal. (C-C″) Control testis stained with Anti-Vasa and Anti-HTS. (D-D″) *Act5C-GAL4 > UAS-CTPsynIsoC* testis stained with Anti-Vasa and Anti-HTS. The branching of fusome occurs relatively early. Scale bar: 20*μ*m.

**Figure 4:**
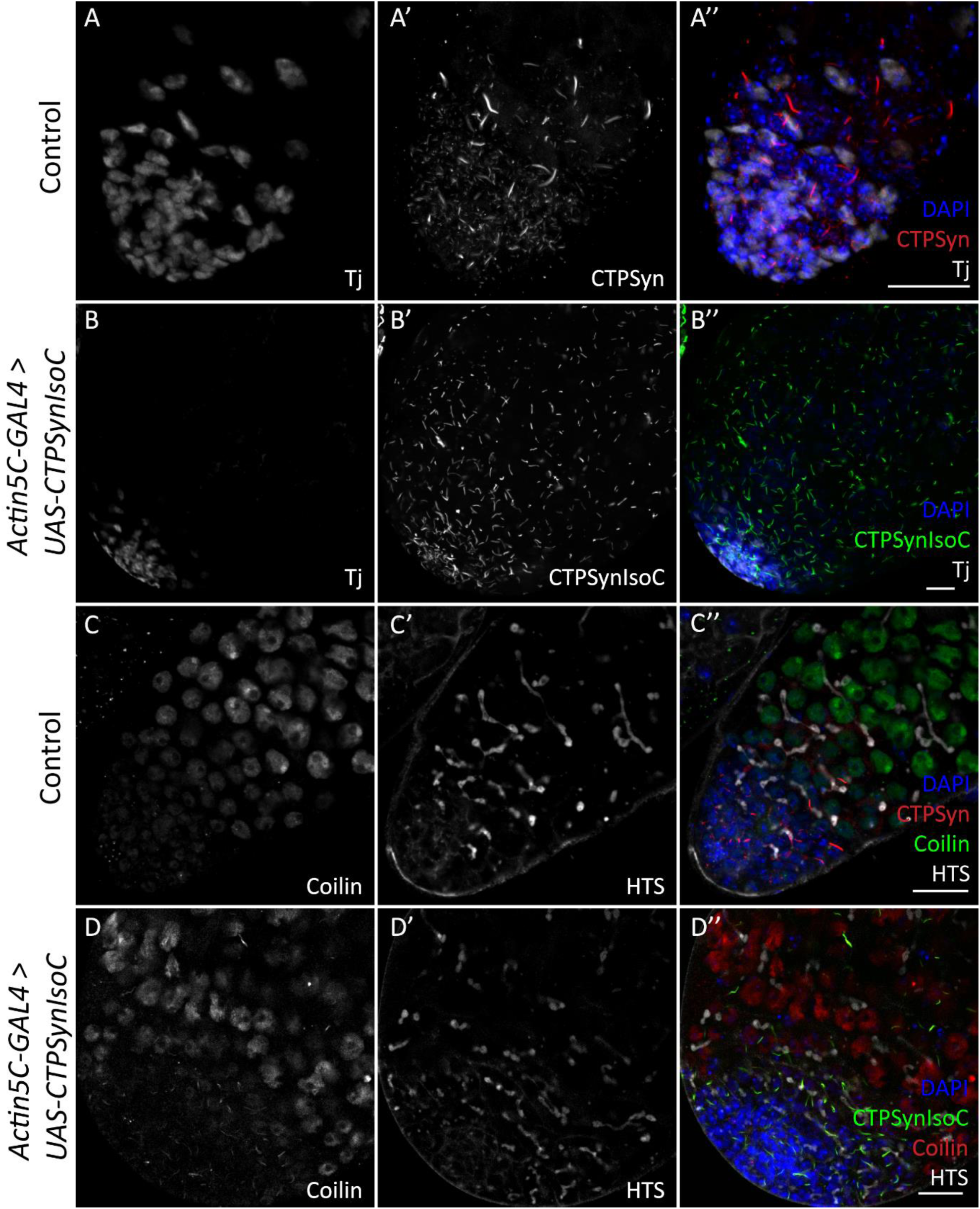
Testes stained with various antibodies. (A – A″) Control testis stained with Anti-Traffic jam and Anti-CTPsyn. The CySC and early cyst cells (Traffic jam) were found to be loosely spaced. (B – B″) *Act5C-GAL4 > UAS-CTPsynIsoC* testis stained with Anti-Traffic jam. The stained cells were more clustered and closely packed. (C – C″) Control testis stained with Anti-Coilin and Anti-HTS. (D – D″) *Act5C-GAL4 > UAS-CTPsynIsoC* testis stained with Anti-Coilin and Anti-HTS. (C – D″) Note that single plane from z-stack images were used instead of maximum projection. Scale bar: 20*μ*m.

**Figure 5:**
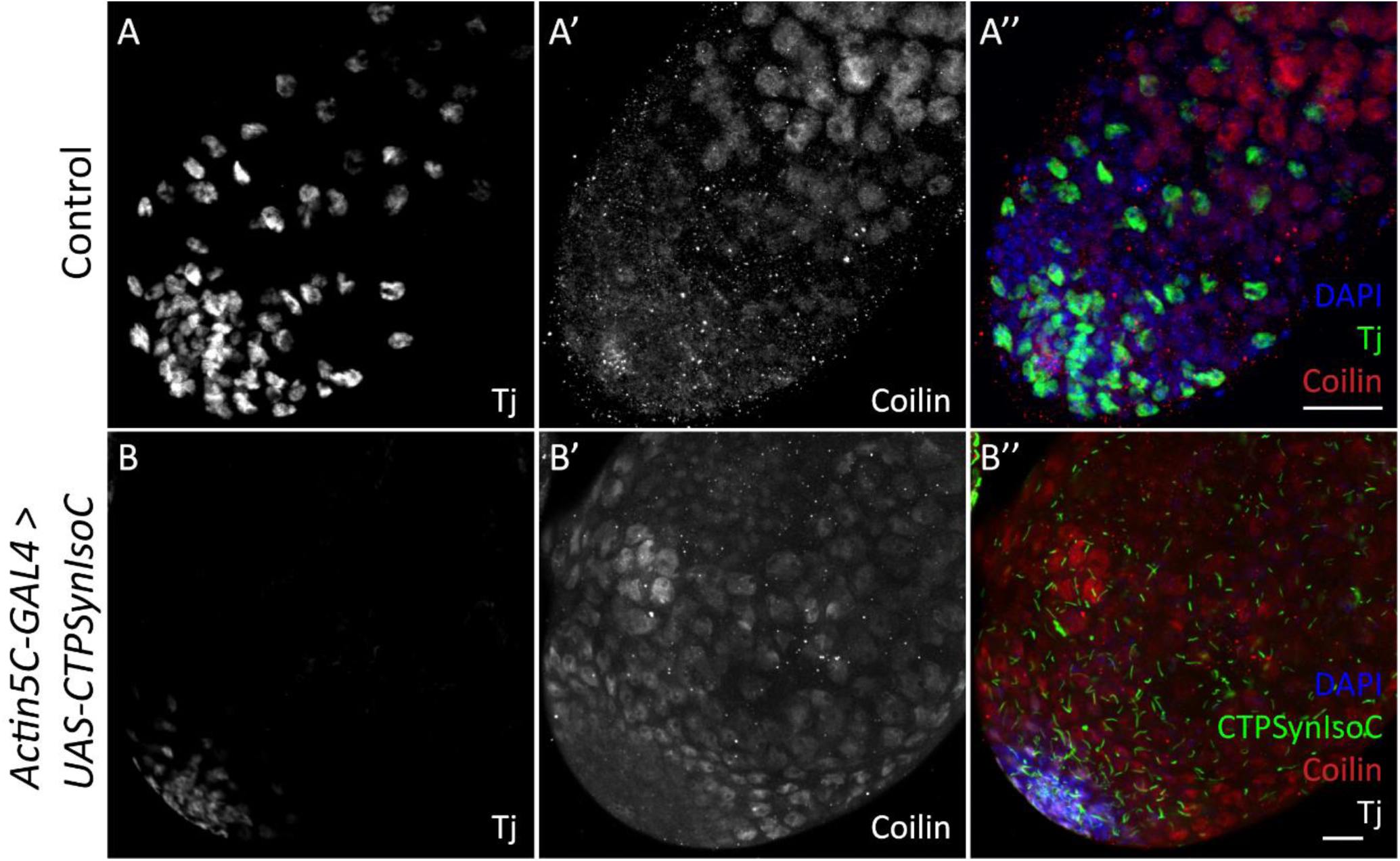
Staining showing testes with Anti-Traffic jam and Anti-Coilin for control testis. (A – A″) and *Act5C-GAL4 > UAS-CTPsynIsoC* testis (B – B″). In control testis, cytoophidia are not found in the later part of the testis when the Traffic jam staining ends [see Figure 4A – 4A″] and Coilin staining starts. Note that (B – B″) is a single plane image instead of maximum projection. Scale bar: 20*μ*m.

We subsequently stained fusome with Anti-HTS (hu-li tai shao) (Lin, Yue, and Spradling, 1994). Fusome is a germline specific organelle. As germline cells develop, the morphology of fusome changes from spherical in GSC to interconnected branches in spermatogonium and its descendants due to incomplete cytokinesis (Hime, Brill, and Fuller, 1996). In control testes, the branching of fusome is observed at a certain distance away from the apical tip. In the *Act5c-GAL4 > UAS-CTPsynIsoC* flies, not only does branching become apparent earlier i.e. in cells closer to the stem cell niche, these structures are found in greater numbers than in controls (Figures 3C′ and 3D′, red arrow). This suggests that germline cells in *CTPsynIsoC-*overexpressing testes not only divide faster, but also divide more, producing larger populations of growing cells. To further document the properties of germline descendants, we used Anti-Coilin to stain for spermatocytes. A GSC divides asymmetrically to give rise to a daughter gonialblast (GB). After four rounds of transit amplification divisions, a spermatogonial cyst comprising of 16 cells is produced. These will later mature into spermatocytes. As Coilin is up-regulated in spermatocytes (Liu *et al*., 2009), an increase in Vasa-positive cell numbers should be accompanied with increased numbers of Coilin-positive cells. We have found this to be true. In control testes, the numbers of Coilin-positive cells across the width of the organ was noted to be slightly less than ten cells, whilst the number increased to approximately 12 - 14 in *Act5c-GAL4 > UAS-CTPsynIsoC* testes (Figures 4C and 4D; also shown previously in Figures 2C and 2D). These findings together show that the ubiquitous overexpression of *CTPsynIsoC* within testicular tissue does indeed induce germline cell proliferation and differentiation, leading to higher cell counts and its bulging phenotype.

### MiR-975 causes similar levels of apical testis bulging and increase of spermatocyte cells in males overexpressing the microRNA

A parallel screening for alterations in cytoophidia phenotypes caused by the overexpression of singular microRNAs (miRNA) was conducted along with our investigation into *CTPsynIsoC* overexpression (Dzaki, Woo, and Azzam, 2018). Lines carrying extra copies of miRNA genes were crossed to various drivers. The morphology of cytoophidia in the ovaries of emergent F1 females was then observed. We saw that flies overexpressing miR-975 displayed significantly lengthier cytoophidia in the nurse cells of their egg chambers, regardless of driver-construct (Figure 6). Cytoophidia within the follicle cells of *T155-GAL4 > UAS-miR975* individuals however, were not affected. Upregulation of *CTPsynIsoC* in ovarian RNA extracts were confirmed via qPCR.

**Figure 6:**
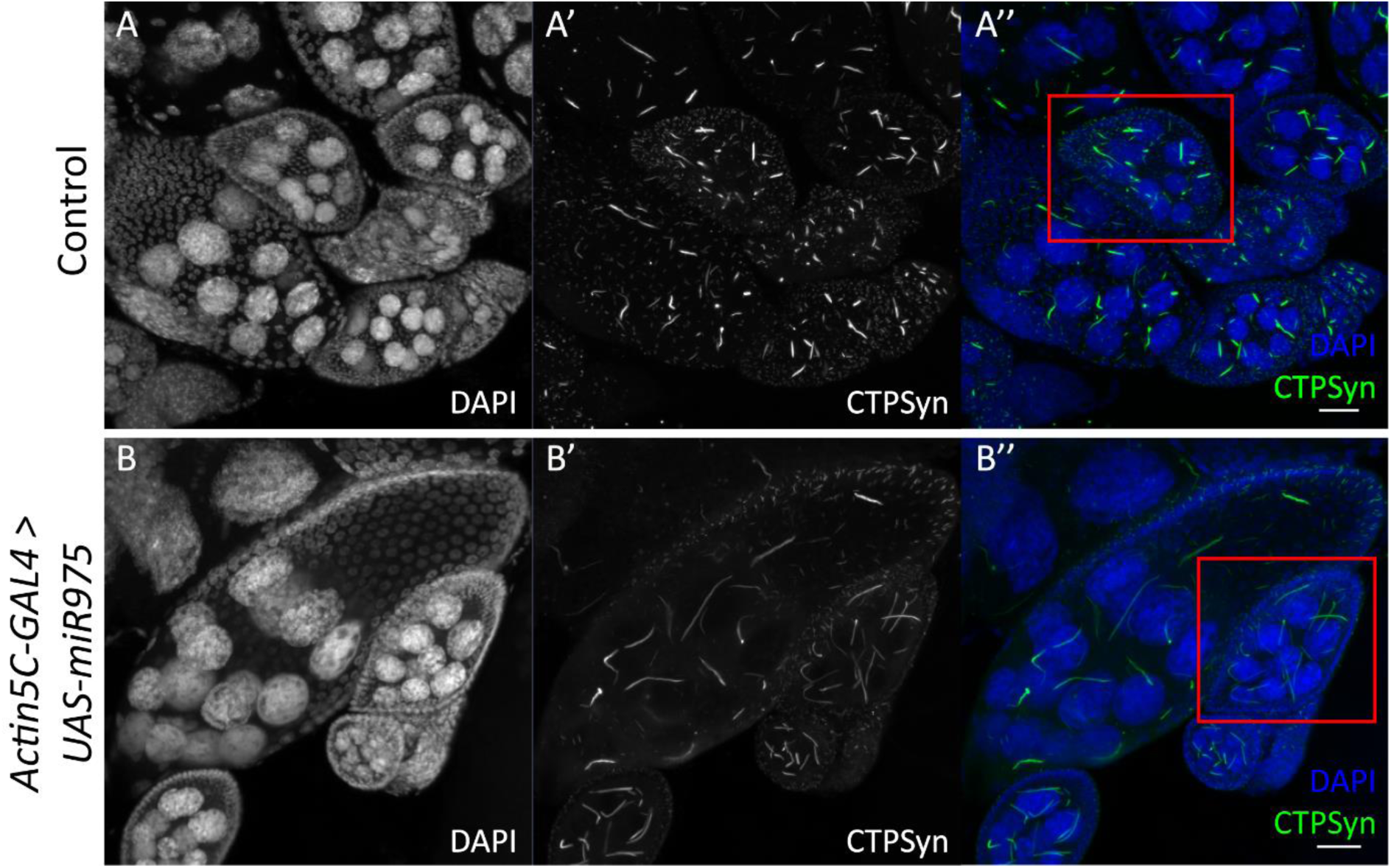
Overexpression of miR-975 induces cytoophidia elongation in egg chambers. (A – A″) Cytoophidia in control ovaries. (B – B″) *Act5C-GAL4 > UAS-miR-975* ovaries. Stage 8 egg chambers are highlighted in red boxes. Note the elongated cytoophidia in B’ and B″. Scale bar: 20*μ*m.

We hypothesized from this information that as *CTPsynIsoC* levels are positively affected by the overexpression of miR-975, like-genotyped males will likely display the same apical bulging phenotype as observed in those overexpressing *CTPsynIsoC*. This was found to be true. Figure 7 clearly shows a circumferential difference between *Act5c-GAL4 > Oregon-R* and *Act5c-GAL4 > UAS-miR975* males. These images further demonstrate a marked increase in fusomic-branching incidences in flies of the latter genotype. Unlike *Act5c-GAL4 > UAS-miR975* females, however, neither cytoophidia length nor numbers appeared affected. Instead, a noticeably greater presence of nuclear CTPsyn – rather than the protein in its tethered form as cytoophidia – was seen beyond the stem cell-niche region in testes tissue overexpressing miR-975. Phenotyping additionally show that these flies would produce similar penetrance levels of bulged testes to that of *Act5c-GAL4 > UAS-CTPsynIsoC* males (Figure 1C).

**Figure 7:**
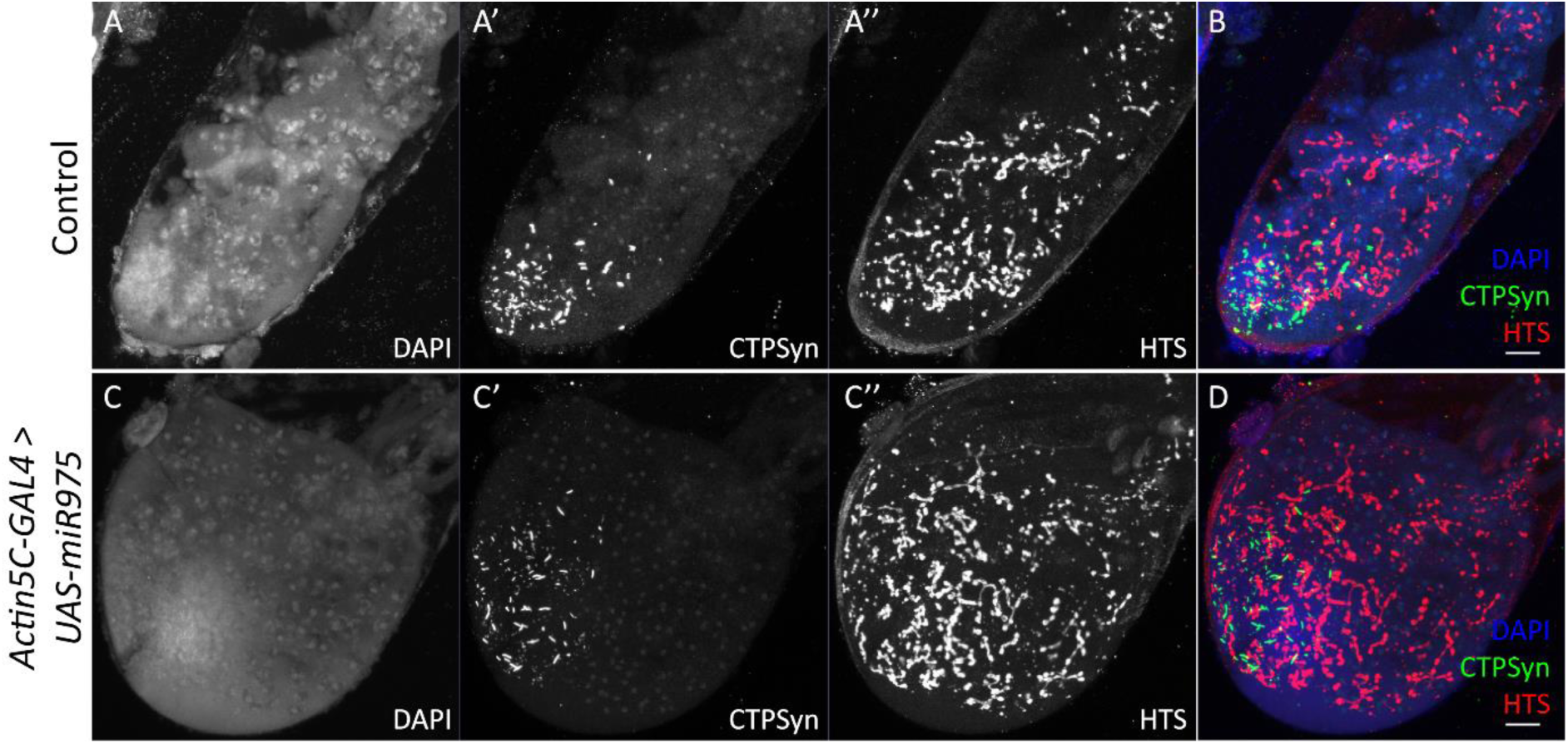
Testes showing CTPsyn and HTS stainings. (A – A″) Control testis, merged shown as (B). (C – C″) *Act5C-GAL4 > UAS-miR975* testis, merged shown as (D). Scale bar: 20*μ*m.

### Overexpression of either *CTPsynIsoC* and *miR-975* lead to upregulated levels of isoforms of *CTPsyn*, but show differential association with commonly-linked genes

Tissue overgrowth is a characteristic hallmark of tumours. It is thus natural to speculate that the bulging testes phenotype observed with either *CTPsynIsoC* or *miR-975* overexpression is attributable to known oncogenic factors. We narrowed a candidate pool down to four genes, all of which have either shown a connection to testicular or prostate cancer in past literature, and/or a demonstrated relationship to CTPsyn (Aughey *et al*., 2016; Fellenberg *et al*., 2010; Mahajan *et al*., 2007). The expression levels of these genes in testes tissue from *Act5c-GAL4 > UAS-CTPsynIsoC* were compared to their expression levels in *Act5c-GAL4 > UAS-miR975*. *Rp49* and *GAPDH* served as references for normalization purposes. Fold-change (fc) values were derived after discounting expressional fluctuations observed in the *Act5c-GAL4 > Oregon-R* controls, where fc is assumed to be zero for each gene of interest (Figure 8).

**Figure 8:**
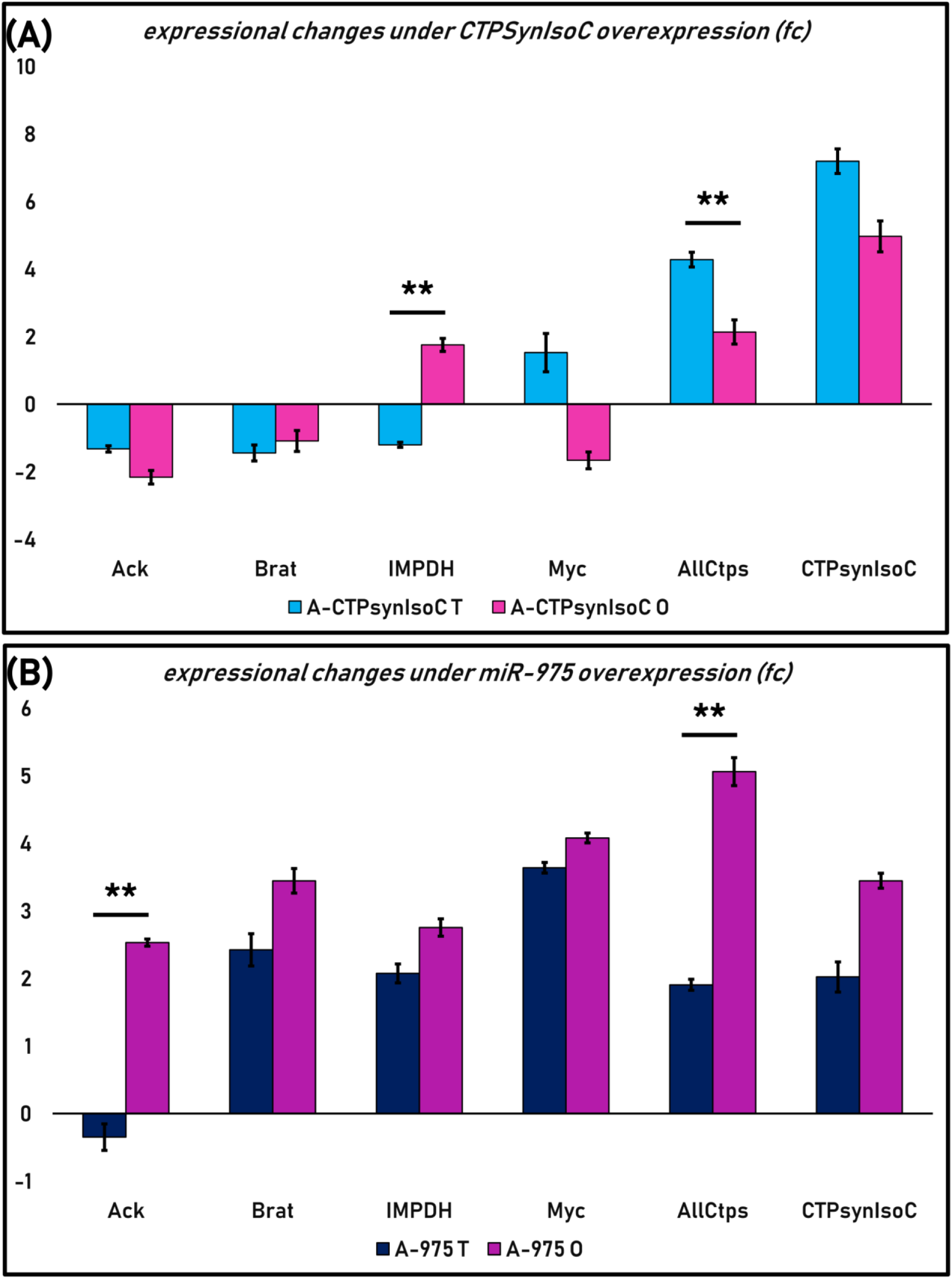
Overexpression of *CTPsynIsoC* and miR-975 each affects expression levels of *CTPsyn*-related oncogenes differently. (A) Overexpression of *CTPsynIsoC* affects *IMPDH* and *Myc* differently in reproductive tissues of different sexes. In (B), miR-975 overexpression only induced such opposite response of the *Ack1* gene. A statistically significant change is marked with ‘**’, where *p*<0.01.

As expected, *Act5c-GAL4 > UAS-CTPsynIsoC* showed a drastic increase in *CTPsynIsoC* levels (fc = 7.20). The primer pair for *AllCtps* was designed to simultaneously capture Isoforms A, B and C of *dme*CTPsyn. A 4.28-fold upregulation of these isoforms was recorded in *Act5c-GAL4 > UAS-CTPsynIsoC* flies. Synergy amongst isoforms is well-documented, and thus this observation was anticipated. However, as we are unable to distinguish isoforms from one another here, the ratios in which each isoform would have contributed to the fold-change would also remain unclear. Nonetheless, as the *Act5c-GAL4 > UAS-CTPsynIsoC* crosses are expected to drive the expression of *CTPsynIsoC* alone, it is likely that the upregulation is attributable to the much higher numbers of this isoform, rather than a positive change in any of the others.

We had expected an upregulation of *CTPsynIsoC* in *Act5c-GAL4 > UAS-miR975* males as well. This was a hypothesis inferred from the cytoophidia-elongating effect that the overexpression of this particular miRNA had on these filamentous structures when they were observed in female egg chambers, and the supporting confirmation that *CTPsynIsoC* was indeed upregulated in ovarian extracts of these flies. We observed moderate upregulations in both *CTPsynIsoC* (fc = 2.02) and *AllCtps* (fc = 1.91). Compared to *Act5c-GAL4 > UAS-CTPsynIsoC* – where the fc disparity between these two were very much obvious – their differences in *Act5c-GAL4 > UAS-miR975* were minute (*p>0.05*). This allows us to deduce that rather than *CTPsynIsoC*, the upregulation of CTPsyn seen in *Act5c-GAL4 > UAS-miR975* testes is attributable in a much more equally-distributed manner amongst the three isoforms.

*Act5c-GAL4 > UAS-CTPsynIsoC* demonstrated downregulations in *Ack, Brat* and *IMPDH*, whilst showing an upregulation in *Myc*. However, fc values were modest across all four genes; none exceeded an fc of 1.6 (fcs = −1.32, −1.44, −1.20, and 1.53, respectively). Fluctuations seen in *Myc* readings between bio-replicates discounts the significance of its upregulation as a causal factor for testes bulging in *Act5c-GAL4 > UAS-CTPsynIsoC* even more. By contrast, *Act5c-GAL4 > UAS-miR975* displayed positive correlations to *Brat, IMPDH* and *Myc* (fcs = 2.42, 2.08 and 3.64, respectively). This indicates that at the very least, excessive levels of the miRNA may be related to the activation of several cancer-related factors, thus leading to the tumour-like phenotype.

## Discussion

### CTPsyn-overexpression potentiates mitotic division within the TA-region

The *Drosophila* testis is a close-ended muscular tube sheathed by pigment cells and actomyosin of genital disc origin (Kozopas, Samos, and Nusse, 1998). In its apical tip, a group of twenty somatic-cells form a region known as the hub. Immediately adjacent to these cells are eight GSCs arranged in a rosette formation, each encapsulated by two CySCs. In its entirety, this unit of functionally-different cells make up the stem cell niche. As asymmetric division occurs during spermatogenesis, ‘core niche members’ retain their non-differentiating features, ensuring that sperm cell production takes place continuously throughout the sexually-mature phase of the lifespan of an adult male fly (Demarco *et al*., 2014).

Paralleled division between GSCs and CySCs gives rise to units of gonialblasts (GBs) surrounded by daughter cyst cells (Hardy *et al*., 1979). Due to the highly-coordinated manner in which this event occurs, cell-to-cell signalling across the niche microenvironment is vital. Somatic and germline cells are known to communicate at the hub through Unpaired-ligand induced activation of JAK-STAT92E (Kiger *et al*., 2001; Tulina and Matunis, 2001). The cooperation between gene products of *bag-of-marbles* (*bam*) and *benign gonial cell neoplasm* (*bgcn*) (Gonczy, Matunis, and DiNardo, 1997) meanwhile regulate differentiation and proliferation of developing spermatogonia within GBs. The activity of cell adhesion protein such as E-cadherin is also required to allow the attachment of both GSCs and CySCs to the hub cells, as well as the maintenance of both the stem cells (Voog, D’Alterio, and Jones, 2008).

Males overexpressing *CTPsynIsoC* have a much greater tendency to produce bulged testes. Tissue overgrowth is always exclusive to the apical tip, where the stem cell niche resides and associated intercellular communication events transpire. In such situations where it is found in excess, is CTPsyn then capable of disturbing the delicate balance observed between these signalling molecules, leading to uncontrolled cell proliferation? Several cancer cell lines of hepatoma (Williams *et al*., 1978), leukaemia (Ellims, Gan, and Medley, 1983; van den Berg *et al*., 1993; Verschuur *et al*., 1998; Whelan *et al*., 1994) and colon (Weber *et al*., 1980) origin are known to intrinsically express higher levels of *CTPsyn*. Conversely, a more recent study has demonstrated how RNAi-mediated knockdown of the gene could reduce tumour overgrowth significantly in *Drosophila* larvae (Willoughby *et al*., 2013). With this study, we ask whether the overexpression of CTPsyn alone could generate tissues or organs with cancerous properties.

In the *Drosophila* testis, each GB undergoes four rounds of transit amplification (TA) divisions. Rapid production of nascent nuclei within such a limited space means that this region outwardly appears dense and tightly-packed under DNA-counterstaining by DAPI. In *Act5c-GAL4 > Oregon-R* controls, greater incidences of micro-cytoophidia is recorded here as well. Beyond the TA region, macro-cytoophidia are typically seen instead. These observations suggest a flexibility in cytoophidia formation, even across relatively small distances. We hypothesize that this interchangeable nature of CTPsyn filamentation might be both a means of regulating the protein’s activity, and a mechanism to modulate response to cellular needs for its CTP product. As aforementioned, the TA region is where active mitotic division occurs continuously. This creates an environment of high-nucleotide demand. The formation of cytoophidium has been shown to signify the inactivity of *dme*CTPsyn (Aughey and Liu, 2015). High CTP demand calls for greater enzymatic activity; more and more CTPsyn proteins thusly dissociate from its filament form, gradually causing a substantial shortening of cytoophidia. CTPsyn is also involved in the synthesis of phospholipids (McDonough *et al*., 1995), another component required during the cytokinesis process following nuclear separation in mitosis.

Vasa and Coilin staining together show that in comparison to controls, *Act5c-GAL4 > UAS-CTPsynIsoC* testes do indeed bear higher numbers of dividing GSCs close to the stem cell niche, accelerating downstream spermatocyte population growth. With its functions so closely tied to mitotic cellular division, it is no wonder then that uncontrolled levels of CTPsyn proteins appear to single-handedly orchestrate over-proliferation events within the TA region, leading to apical tip bulging.

### Testes as a model for studying overgrowth-typed germline cell tumours in *Drosophila*

*CTPsynISoC-*overexpression induced lengthening of cytoophidia was first confirmed by immunohistochemistry of ovaries. Apart from abnormally long cytoophidia and a minor shrinkage in terms of size, however, no other side effects from the resultant excess in its protein levels were observed within these tissues. Human *CTP synthase 1* i.e. *CTPS1* upregulation has long been implicated in cancer (Williams *et al*., 1978). As *CTPsynIsoC* is the direct orthologue of *hsaCTPS1* for *Drosophila melanogaster*, we had thus expected to see hallmarks of the disease within cells overexpressing the gene. These should be driver-dependent: manifestation would appear as either germline cell tumours (GCTs) under *Act5c-GAL4*, or irregular numbers of cysts within developing egg chambers under *nanos-GAL4* (Forbes and Lehmann, 1998; Salz, Dawson, and Heaney, 2017). Repeated attempts unfortunately show GCT counts in flies overexpressing *CTPsynIsoC* to be only comparable to controls, and that the number of cysts displaying anything beyond the normal 15:1 nurse cell to oocyte ratios have remained non-significantly changed.

To further explore cancer in regards to *CTPsynIsoC* overexpression, we therewith shifted our focus onto the fly testis. Whilst testes GCTs are thus far under-described for *Drosophila*, some of the underlying causes of ovarian GCTs appear to have a ‘male’ context. For example, the loss of the female specific *sex-lethal* (*Sxl*) in adult females induces an upregulation of male genes (Chau, Kulnane, and Salz, 2009) such as *PHD-finger-7* (*phf7*), a known regulator of male germ cell fate during spermatogenesis (Yang, Baxter, and Van Doren, 2012). Build-up of male-specific transcripts leads to an inevitable loss in female identity, negatively affecting differentiation and eventually producing the GCT phenotype (Shapiro-Kulnane, Smolko, and Salz, 2015). This in itself is arguably ‘masculine’; the progression of cystoblasts in ovarian GCTs very closely mimics how spermatogonial cysts give rise to spermatocytes. It is thus quite astonishing why *Drosophila* testes GCTs is as poorly defined as it is: the inherent maleness of female-manifested tumours, as well as recent evidence providing proof that *hsaCTPS1* is intensely upregulated in testicular cancer tissue (Uhlen and Zhang, 2017) in our opinion only makes testes an even more attractive prime study material for cancer-related studies.

More than 30% of *CTPsynIsoC*-overexpressing males presented with overgrown testes. Bulging is often restricted to the apical region. We have already discussed the cell populations which likely contributed to the phenotype based off of immunostaining outcomes. The primary cause of bulging may have been the rapid lateral division of GSCs into gonialblasts (GBs). Whereas GBs would detach physically away from GSCs under normal growth conditions, this was not the case where *CTPsynIsoC* was expressed excessively. Subsequent TA-events within developing cysts therefore occur outwardly, rather than down the ‵neck of the testes as it should be. The resulting expansion-of-girth phenotype is what we now categorically recognize as ‘bulging’.

Elongated cytoophidia typical of *CTPsynIsoC* overexpression are found indiscriminately within this region and beyond. No cell type appeared free of the structure. One potentially important observation is their conserved lengths. As aforementioned, cytoophidia are abundant in control testes as well. However, spatial and temporal separation between occurrences of micro and macro-cytoophidia are commonly observed. These incidences are thought to coincide with division activity as a mechanism of adaptive metabolic regulation (Aughey *et al*., 2014; Aughey and Liu, 2015; Strochlic *et al*., 2014). In *Act-GAL4 > UAS-CTPsynIsoC* testes, no such distinction is apparent. Cytoophidia seem to instead be of uniform size and length in all cells where they are observed, regardless of the stage of spermatogenesis progression the cell represents. This indicates that there may be limitations in the extent to which CTPsyn could tether to each other in male testes. *Drosophila* cytoophidia is likely heterogenous (Liu, 2011), especially in germline cells (Liu, 2010). Where CTPsyn protein levels are as high as they are expected to be in *Act-GAL4 > UAS-CTPsynIsoC*, further oligomerization may thusly be limited by the availability of its hypothetical associated factor(s). We have already elaborated upon the critical roles of CTPsyn as a metabolic enzyme. Excessive CTPsyn activity is known to encourage nucleic acid replication and proliferation (Martin *et al*., 2014). Its involvement in membrane phospholipid production additionally implicates CTPsyn in cellular division events which naturally follow immediately after (Chang and Carman, 2008). As cytoophidia formation literally falls short, we believe these synergistic consequences of heightened CTPsyn activity are what may have disrupted normal cell growth, culminating in the tumour-like bulging phenotype.

Observations in *CTPsynIsoC*-overexpressing ovaries might provide further support to this purported relationship between cytoophidia length and tumorigenesis (Azzam and Liu, 2013). These filaments are remarkably long in both somatic and germline cells of the tissue, even in comparison to testes cytoophidia in males of the same genotype. Subsequent screening for signs of GCTs and unordinary cyst cell growth patterns came out negative. Is there then a cause and consequence correlation between cytoophidia length, metabolic dysregulation, and tumour-like growth? As there are stark contrasts in how the germline cell responds to over-production of CTPsyn, another interesting arising question is whether sex-specific factors are involved; if so, which mechanisms involving CTPsyn are they influencing, and how does this lead to the development of testes GCTs? In the practice of utilizing *Drosophila* to study cancer, its brain is most commonly used as a model, especially in regards to tumour overgrowth (Beaucher *et al*., 2007; Sonoda and Wharton, 2001). Certain germline genes have in fact been ectopically expressed in this tissue in order to demonstrate their malignant-growth inducing abilities (Janic *et al*., 2010). An acknowledged disadvantage of ectopic expression is that outcomes may not accurately reflect *in situ* outcomes. This might be even more critical when genes transverse germ to soma; micro-environmental influence is thought to be especially instrumental in the development of human GCTs (Diez-Torre *et al*., 2010; Luo *et al*., 2016). We therefore reiterate the massive potential of *Drosophila* testes as a model of GCT-study, particularly where the cause of GCTs is sex-specific, and pathological expectations include visible overgrowth.

### Overgrowth of testes observed in both overexpression lines is associated with hyper-production of CTPsyn, but is produced through diverging mechanisms

Similar to *CTPsynIsoC*-overexpression, initial screening phases into identifying cytoophidia-altering miRNA overexpression events saw us exclusively scrutinizing ovarian cells as well. Our choice to do so was also in part due to limitations imposed by expressional characteristics of employed GAL4-drivers. For most miRNAs, we were doubly limited by the purported inadequacy of the UASt construct within germline cells (Rørth, 1998). Regardless, concurrent utilization of both follicle-cell and nurse-cell capable drivers allowed us to effectively isolate a small group of candidate miRNAs by their effects on cytoophidia morphology (Hudson and Cooley, 2014).

Our miRNA-overexpression screen revealed that when expressed in excess, miR-975 asserts an elongating effect upon ovarian cytoophidia. We now know that observed elongation in females could translate into testicular apical tip enlargement in a third of emergent males. Though less extreme than the *Act-GAL4 > UAS-CTPsynIsoC* phenotype, the lengthening of cytoophidia was still prominent enough in *Act5c-GAL4 > UAS-miR975* females for us to be optimistic about obtaining bulged testes in like-genotyped males. RT-qPCR further confirmed a modest upregulation of *CTPsynIsoC* in testes overexpressing *miR-975*.

Subsequent phenotyping revealed that not only are *Act5c-GAL4 > UAS-miR975* males producing bulged testes, but a significantly high proportion of them do (20.56%; n=224). Of course, the relationship between MicroRNAs and cancer is well studied (Jansson and Lund, 2012). The critical roles they play in monitoring gene expression often mean that with their dysregulation, the disruption to cellular ‘normal’ which typically follow may eventually lead to cancerous phenotypes. Many miRNAs mimic tumour suppressors in functionality (Sassen, Miska, and Caldas, 2008). An inverse relationship with carcinogenesis is thusly expected in most cases. Several miRNAs are nonetheless known to encourage the progression of certain cancer types such as lung and hepatocellular carcinoma, rather than repressing them (Su *et al*., 2014; Wu *et al*., 2015). The trio-clustered *hsa*miRNAs 371, 372, and 373 have proven themselves to be potent oncogenes. Their part in driving testicular germ cell tumours in particular is widely acknowledged, so much so that measuring sera *hsa-miR-371* levels is now applied as a diagnostic biomarker tool (Dieckmann *et al*., 2012).

In this study, the overexpression of *miR-975* alone appears to have sufficiently induced visible tissue overgrowth in *Drosophila* testes. As with *CTPsynIsoC*-overexpressing flies, we endeavoured to understand how this phenotype could have been caused by staining bulged testes with anti-HTS. The primary antibody stains for Adducin-like proteins, found within fusomes. These proteins are divided between two nascent fusome organelles during cell-division. However, due to the incomplete nature of cytokinesis amongst dividing spermatogonial cells, fusomic-division is by consequence also incomplete; Adducin-like proteins are thus highlighted as thin structures inter-bridging neighbouring spermatogonia within developing cysts. These are referred to as ‘fusomic-branches’ (Matunis, Stine, and de Cuevas, 2012). Figure 7 demonstrates that in comparison to control, *miR-975-* overexpressing testes show higher incidences of fusomic-branching. This overlaps patterns seen with anti-HTS staining in *CTPsynIsoC-*overexpressing males. As fusomes are temporally seen as early as within gonialblasts (GB), we can conclude that akin to *Act-GAL4 > UAS-CTPsynIsoC*, testes bulging in *Act-GAL4 > UAS-miR975* too is attributable to excessively robust GB production around the stem cell niche. Girth-expansion may then be a result of the tissue accommodating above-normal spermatocyte population sizes.

Although they share these key traits, several differences between *Act-GAL4 > UAS-CTPsynIsoC* and *Act-GAL4 > UAS-miR975* must be noted. For one, cytoophidia is not elongated in *miR975*-overexpressing testes. Based on the established association between cytoophidia and testicular bulging in *CTPsynIsoC-*overexpressing flies as well as preliminary RT-qPCR data, we had anticipated these tissues to bear much lengthened filaments. Rather, they appeared unaltered, with lengths and numbers comparable to that of control testes (*Act-GAL4 > Oregon-R)*. As a matter of fact, the opposite may instead be true. Cytoophidia presented in *miR-975*-overexpressing testes outwardly appears less compacted than even the controls. These observations nonetheless uphold the theory of correlation between control of cytoophidia formation and internal metabolism (Aughey *et al*., 2014; Aughey and Liu, 2015). Though higher levels of *CTPsynIsoC* are seen in *Act-GAL4 > UAS-miR975* flies, fc values suggest that upregulation is modest. This increment in *CTPsyn* may be insignificant for counterbalance mechanisms such as filamentation into cytoophidia to be kick-started into action. With its protein levels left unchecked, the same lateral growth of cells are encouraged, leading to the bulged phenotype. We must also press that whereas the structure was found throughout the testicular body in *Act-GAL4 > UAS-CTPsynIsoC* males, it remained contained within the boundaries of the TA-region under *miR-975*-overexpression. An enrichment of nuclear-*CTPsyn* was however obviously detectable in downstream cysts. This trait of *Act-GAL4 > UAS-miR975* bulged testes along with its under-compacted cytoophidia morphology has led us to believe that *miR-975* may regulate a physical constituent of cytoophidia formation. Its downregulation as a consequence of *miR-975* overexpression eventually restricts CTPsyn tethering. A more optimistic extension to the theory is that this physical constituent is unique to male germline cells, as cytoophidia elongation occurred freely in like-genotyped females. Co-precipitation analysis as well as transcriptomics approaches are currently underway to explore these and the many other possibilities further.

It is already apparent that whilst overexpression of *CTPsynIsoC* and *miR-975* both culminates in a similar tumour-like overgrowth, the pathways involved in either case may be completely divergent of one another. There is demonstrable proof of the oncogene-like effects of *CTPsyn* in mammalian tissue. With this study, we have shown that when overexpressed, the gene bears oncogene-like effects upon *Drosophila* tissue as well. A conservation of function amongst gene orthologues is a fact of biology (Tatusov, Koonin, and Lipman, 1997). It is not a wonder then that, even in two vastly different organisms, *CTPsyn* would assert the same consequences. As aforementioned, a cluster of *hsa-*miRNAs come highly implicated in carcinogenesis of testicular tissue (Voorhoeve *et al*., 2006). To our misfortune, not only do *Drosophila* appear to not carry their orthologues, but *miR-975* may also be genus specific: on miRBase, the miRNA has only been reported for several drosophilid specie (Kozomara and Griffiths-Jones, 2011). This means that information regarding potential functions of this small-RNA is scarce, although *in silico* target prediction (Kheradpour *et al*., 2007) has shown us that many of the mRNAs it may elicit are involved in membrane-trafficking, cell-cycle regulation and DNA-repair. Due to study constraints, we are unable to transcriptomically verify the validity of these predictions. A much narrowed approach to understanding *miR-975* involvement in the advent of cancer within *Drosophila* tissues was therefore attempted through RT-qPCR.

We have already discussed how *CTPsyn* may be connected to *Ack1* (Strochlic *et al*., 2014), *IMPDH* (Chang *et al*., 2015), and *Myc* (Aughey *et al*., 2016), either genetically or proteomically. *Brat* was included as a well-known translational repressor of brain tumours in *Drosophila* (Sonoda and Wharton, 2001). Given the nature of their reported relationships, all genes apart from *Brat* should be upregulated in *Act-GAL4 > UAS-CTPsynIsoC* testes. RT-qPCR results instead deviated from expectations as both *IMPDH* and *Ack1* were downregulated. However, any changes observed were, in our opinion, too mild to be of significance. None of the fc values exceeded 1.5. By contrast, expressional changes seen with *Act-GAL4 > UAS-miR975* testes were more drastic. The overexpression of *miR-975* is accompanied by much higher *Myc* and *IMPDH* levels. As *Myc-*lacking follicle cells have been shown to be completely lacking in cytoophidia (Aughey *et al*., 2016), its upregulation must have therefore contributed to the concurrently-observed upregulation of *CTPsynIsoC* in *Act-GAL4 > UAS-miR975* flies. Well-documented functions of *Myc* as a potent oncogene may have also led to excessive proliferation and evasion of apoptosis, propagating tumorous tissue growth (Dang, 2012).

The relationship between *CTPsynIsoC* and *IMPDH* appears to be more complex. IMPDH is another protein which compartmentalizes into cytoophidia as a means of activity control (Hedstrom, 2009). It is able to form cytoophidium both together (Carcamo *et al*., 2011) and independently of CTPsyn (Chang *et al*., 2015). A point of note is that whilst both genes are upregulated to similar degrees in male and female *Act-GAL4 > UAS-miR975* RNA extracts, cytoophidia elongation was not observed in testes tissue. Increased IMPDH-filamentation has been shown to encourage CTPsyn-filamentation (Chang *et al*., 2015), and vice versa. Chang’s study further reported that under conditions whereby *CTPsyn* upregulation is causal to greater IMPDH-filamentation, the event takes place despite cellular IMPDH levels remaining unchanged. Our RT-qPCR data instead suggest that in *miR-975* flies, *IMPDH* gene expression is induced alongside *CTPsyn*. We suspect that rather than being affected directly by *CTPsyn*, however, it is the drastically higher levels of *Myc* which is responsible for *IMPDH* upregulation, as the latter has been known to regulate nucleotide biosynthesis through IMPDH-activity (Liu *et al*., 2008). In the context of how IMPDH and CTPsyn could have co-regulated one another’s compartmentation behaviour, it appears that the mechanisms at play here may also be subject to sex-specific factors.

What weakens our stance for promoting *miR-975* as an onco-miRNA is how *Brat* expression levels was also positively affected by its overexpression, where we had conversely expected its downregulation. Functionally, Brat is powerful enough to markedly decrease *dme*Myc protein levels in various *Drosophila* tissues (Gallant, 2013). We can only speculate that the compounding effects of *Myc* and *IMPDH* upregulation, as positive drivers of cell growth, trumped any down-regulatory effects *Brat* may have as a tumour suppressor. We have also reiterated the importance of considering the possibility of sex-disparate phenotypes in GSCs time and time again within this study: whereas the relationship between Brat and *dme*Myc has been demonstrated in that of female’s (Neumüller *et al*., 2008), it has yet to be identified within that of male’s. Further investigations involving detailed microarray data comparison between female and male GSCs should provide a more in-depth understanding.

## Conclusions

In this study, we demonstrate that *CTPsynIsoC* overexpression causes a lengthening of cytoophidia in testes tissue. For a third of males, this culminates in the form of apical tip bulging. Higher numbers of germline cells and spermatocytes is shown to be main cause of testicular overgrowth, as these are stained by Anti-Vasa and Anti-Coilin, respectively. A microRNA i.e. miR-975 is also found to induce the bulged-testes phenotype in ∼20% of males overexpressing it, although cytoophidia length, size, and compaction is not visibly altered. RT-qPCR nonetheless reveals that a small but significant increment in *CTPsynIsoC* levels does accompany *miR-975* overexpression. This is not a two-way relationship, *CTPsynIsoC* overexpression does not cause *miR-975* upregulation. Quantification of a number of cancer-related genes further show how differentially either overexpression events affect genetic expression, suggesting a divergence in the pathways they potentially regulate. They both do, however, affect *Myc* expression positively. As a potent oncogene, it is possible that the defining induction event in testes overexpressing either *CTPsynIsoC* or *miR-975* is Myc-protein production, the effects of which would have eventually led to excessive proliferation, and therefore tissue-overgrowth. Though the findings presented here are insufficient to solidly link *CTPsynIsoC* or *miR-975* to tumourigenesis, this study has demonstrated that the overexpression of either gene could affect tissue growth in such a way that it culminates in a phenotype akin to tumours. For this reason, we also propose greater utilization of testes as a source of GCT studies. In the future, simultaneous overexpression of *CTPsynIsoC* and *miR-975* could be employed to further investigate if there is a connection between the enzyme and the miRNA. Knock-down experiments using RNAi and transcriptomics as well as RNA sequencing will also be implemented to identify the causes of the bulging phenotype, and to clarify once and for all whether these underlying mechanisms are indeed cancerous by nature.

## Materials and Methods

### Fly stocks

Stocks were raised at 25°C on cornmeal; the recipe has been slightly modified to suit the local climate and availability of ingredients. The sole driver-GAL4 line involved in this study is *Act5C-GAL4/CyO* (Kyoto *Drosophila* Genome and Genetic Resources [KGGR] #107727). Progenies of *Act5C-GAL4/CyO* crossed to Oregon-R flies (we refer this line as *Act5C-GAL4 > Oregon-R* in this article) were controls for all experiments unless stated otherwise (Kramer and Staveley, 2003; Rezaval, Werbajh, and Ceriani, 2007). The overexpression line i.e. *UAS-UAS-CTPsynIsoC* was a gift from the Liu lab in DPAG, University of Oxford (Azzam and Liu, 2013). The expression of this construct produces Venus-tagged CTPsynIsoC proteins. Crosses made were grown at 28°C in a 12h/12h light to dark cycle.

### Immunohistochemistry

F1 flies were flipped onto wet yeast. Males aged 5 to 6 DAE were dissected in ice-cold 0.2% Triton X-100 in 1X PBS (PBST) in quick succession. Testes were fixed in freshly-made 4% paraformaldehyde (pH 7.4) in 0.5% Triton X-100 in 1X PBS for 30 minutes at room temperature. Successive washing in PBSTX (0.3% Triton X-100 + 0.1% Tween-20 in 1X PBS) and blocking steps were adopted from a previously detailed protocol (Kibanov, Kotov, and Olenina, 2013). Primary antibodies used in this study were rabbit anti-Vasa (1:2000, R. Lehmann), rabbit anti-CTPsyn (1:200, J.L. Liu), guinea pig anti-Traffic Jam (Tj) (1:5000, D. Godt), rabbit anti-Coilin (1:2000, J.L. Liu), guinea pig anti-Coilin (1:2000, J.L. Liu), mouse anti-Fasciclin III (1:40, Developmental Studies Hybridoma Bank [DSHB]), and mouse anti-1B1 (or hu-li tai shao [HTS]) (1:200, DSHB). Tissues were stained in primary antibody solutions overnight at room temperature. Secondary antibodies used were goat anti-rabbit DyLight 488, goat anti-rabbit DyLight 550 (1:600), goat anti-mouse DyLight 550 (1:600), goat anti-mouse DyLight 633, goat anti-guinea pig DyLight 488, and goat anti-guinea pig DyLight 633. All were purchased from Invitrogen and used at 1:400 dilutions, unless stated otherwise. After overnight nutation at room temperature, tissues were mounted with SlowFade Diamond Antifade Mountant (Thermo Fisher Scientific, USA, Cat. No. S36972).

### Microscopy

Images were acquired under 10X, 20X, 40X and 63X lenses on the LSM 710 confocal microscope platform (Zeiss, Oberkochen, Germany) with manufacturer-supplied software, Zen Blue (2012 Edition). Standard objectives were maintained, unless specified.

### Phenotyping assay

Remaining males from crosses described above were dissected in batches of fifty. Testes were classified on the basis of apical tip size. A phenotypically significant testis was scored if the apical tip was bulging; in the case where the tip was not bulged, an increase in size in the neck was considered instead. Statistical tests were conducted by presenting raw data into Excel sheets (Microsoft, CA). Two-way Student T-Tests was performed. Error bars represent the standard error of sum values. A *p-*value below 0.05 indicated significant deviations from controls.

### RNA extraction

Testes were dissected out into double-sterilized PBST within certain time constraints. A maximum of thirty males were processed per batch. PBST was replaced with ice-cold TRIzol® (Invitrogen, USA) and testes samples were immediately frozen at −20°C. Three batches were consolidated prior to homogenization by QiaShredder spin columns. Subsequent RNA extraction was achieved using the RNeasy® Mini kit (all by Qiagen, Hilden, Germany). All extracts underwent DNase digestion (RNase-free DNase set, Qiagen Biotechnology, USA) prior to quantification.

### cDNA synthesis and qPCR

Complementary-DNA (cDNA) was synthesized with iScript™ Reverse Transcription Supermix (Bio-Rad, USA) according to manufacturer’s instructions. Primers utilized in quantitative-PCR (qPCR) are listed in Supplementary Table S1. Reactions were prepared in 10*μ*l volumes with iTaq™ Universal SYBR® Green Supermix before being loaded onto the CFX96 qPCR platform (all from Bio-Rad, Hercules, California, USA). Standard qPCR protocol for all plates entailed an initial denaturation of 2 minutes at 95°C, followed by 40 cycles of fluorescence-quantification with denaturation, annealing and extension at 95°C/10s, 59°C/15s and 72°C/15s, respectively. Melting-curve analysis with temperatures between 65°C to 95°C succeeded every run. Data analysis was performed manually on Excel (Microsoft, California, USA) with aid from Bio-Rad’s own software where possible.

---

**Supplementary Table S1:**
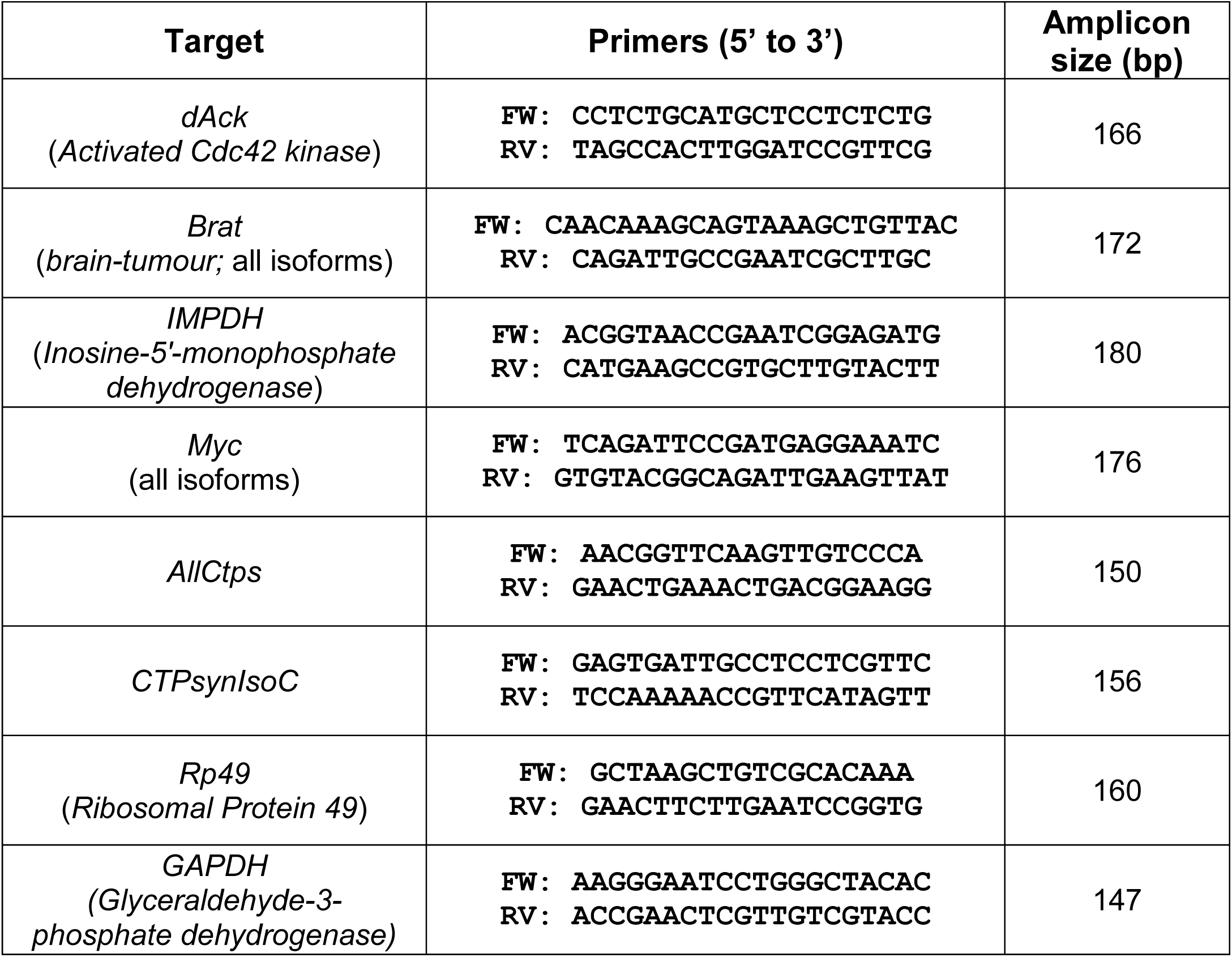
Primers for Qpcr

